# Molecular characterization and differential expression reveal functional divergence of stress-responsive enzymes in C_4_ panicoid models, *Setaria italica* and *Setaria viridis*

**DOI:** 10.1101/2019.12.24.887927

**Authors:** Mehanathan Muthamilarasan, Roshan Kumar Singh, Bonthala Venkata Suresh, Priya Dulani, Nagendra Kumar Singh, Manoj Prasad

## Abstract

Stress-responsive genes regulate the morpho-physiological as well as molecular responses of plants to environmental cues. In addition to known genes, there are several unknown genes underlying stress-responsive machinery. One such machinery is the sophisticated biochemical carbon-concentrating mechanism of the C_4_ photosynthetic pathway that enables the plants to survive in high temperatures, high light intensities and drought conditions. Despite the importance of C_4_ photosynthesis, no comprehensive study has been performed to identify and characterize the key enzymes involved in this process among sequenced Poaceae genomes. In the present study, five major classes of enzymes that are reported to play roles in C_4_ biochemical carbon-concentrating mechanism were identified in sequenced Poaceae genomes with emphasis on the model crops, *Setaria italica* and *S. viridis*. Further analysis revealed that segmental and tandem duplications have contributed to the expansion of these gene families. Comparative genome mapping and molecular dating provided insights into their duplication and divergence in the course of evolution. Expression profiling of candidate genes in contrasting *S. italica* cultivars subjected to abiotic stresses and hormone treatments showed distinct stress-specific upregulation of *SiαCaH1*, *SiβCaH5, SiPEPC2*, *SiPPDK2*, *SiMDH8* and *SiNADP-ME5* in the tolerant cultivar. Altogether, the study highlights key stress-responsive genes that could serve as potential candidates for elucidating their precise roles in stress tolerance.

**Key message:** Comprehensive analysis of stress-responsive gene families in C_4_ model plants, *Setaria italica* and *S. viridis* identified *SiαCaH1*, *SiPEPC2*, *SiPPDK2*, *SiMDH8* and *SiNADP-ME5* as potential candidates for engineering abiotic stress tolerance.

## Introduction

Photosynthesis converts light energy into chemical energy to fix atmospheric carbon dioxide (CO_2_) for synthesizing reduced carbon compounds. Three distinct mechanisms of carbon fixation, namely, C_3_, C_4_ and crassulacean acid metabolism (CAM) have been reported in terrestrial plants (Furbank and Taylor 1995). In C_3_ photosynthesis, plants assimilate CO_2_ through the Calvin cycle that occurs inside of the chloroplast in the mesophyll cells (Furbank and Taylor 1995). In C_4_ plants, CO_2_ is converted into bicarbonate followed by its carboxylation to form a four-carbon compound, oxaloacetate (OAA) in the mesophyll. OAA undergoes several possible modifications and diffuses into bundle sheath, where decarboxylation and refixation of CO_2_ by Rubisco in the Calvin cycle takes place (Furbank and Taylor 1995). The mesophyll-located C_4_ cycle serves as a biochemical CO_2_ pump to elevate the CO_2_ levels in the bundle sheath and ensures an uninterrupted supply of CO_2_ to Rubisco, thus enhancing the photosynthetic water use efficiency of C_4_ plants (Schuler et al. 2016). Also, Nitrogen requirement in the C_4_ leaves is lower due to a higher Rubisco catalytic turnover rate, which confers better photosynthetic nitrogen use efficiency in these plants (Schuler et al. 2016). C_4_ photosynthesis is common among monocots, but not very common among dicots (Furbank and Taylor 1995; Schuler et al. 2016). Particularly, the grass family, Poaceae constitutes C_3_, C_3_-C_4_ intermediate and C_4_ species (Grass Phylogeny Working Group II 2012). Among C_3_ grasses, rice (Erhartoideae) and wheat (Pooideae) are considered as staple crops, whereas barley and Brachypodium (Pooideae) are equally important as bioenergy feedstocks. The PACMAD (Panicoideae, Arundinoideae, Chloridoideae, Micrairoideae, Aristidoideae and Danthonioideae) clade of Poaceae is C_4_-rich (Grass Phylogeny Working Group II 2012), where the important C_4_ grasses including sorghum, maize, millets and sugarcane are present in the Panicoid tribe.

Among the Panicoids, *Setaria italica* (foxtail millet) and its wild progenitor, *S. viridis* (green foxtail) are considered as model crops for studying C_4_ photosynthesis (Brutnell et al. 2010; Li and Brutnell 2011), abiotic stress tolerance (Diao et al. 2014; Muthamilarasan and Prasad 2015; Huang et al. 2016), biofuel (Lata et al. 2013; Bandyopadhyay et al. 2017) and nutritional traits (Muthamilarasan et al. 2016; Pant et al. 2016). These crops (collectively termed *Setaria*) possess small diploid genome (~500 Mb), self-pollination and inbreeding nature, shorter life-span (~90 days; seed-to-seed), short stature (~100-150 cm average) and prolific seed production (several thousand seeds per panicle; Lata et al. 2013). The genomes of *S. italica* and *S. viridis* have also been sequenced (Bennetzen et al. 2012; Zhang et al. 2012), which has enabled the development of several genetic and genomic resources in *Setaria* (Muthamilarasan and Prasad 2015). In structural genomics perspective, large-scale, genome-wide molecular markers were developed (Pandey et al. 2013; Kumari et al. 2013; Muthamilarasan et al. 2014a; Zhang et al. 2014) and high-density genetic linkage maps were constructed for mapping the genomic regions controlling agronomic traits (Jia et al. 2013; Fang et al. 2016). In functional genomics, several important gene families including NAC transcription factors (Puranik et al. 2013), WD40 proteins (Mishra et al. 2014), MYB transcription factors (Muthamilarasan et al. 2014b), AP2/ERF transcription factors (Lata et al. 2014), C_2_H_2_-type zinc finger proteins (Muthamilarasan et al. 2014c), Nuclear Factor Y (Feng et al. 2015), 14-3-3 proteins (Kumar et al. 2015), WRKY transcription factors (Muthamilarasan et al. 2015), heat-shock proteins (Singh et al. 2016), SET domain-containing proteins (Yadav et al. 2016), autophagy-associated genes (Li et al. 2016), pentatricopeptide repeat proteins (Liu et al. 2016), DOF containing proteins (Zhang et al. 2017) and HD-Zip transcription factors (Chai et al. 2018) were identified and characterized. However, no elaborate study on C_4_ photosynthetic genes has been performed in any Panicoid species including *Setaria* to understand the structure, organization, evolution and expression profiles in response to environmental perturbations.

In the present study, five major classes of enzymes that play pivotal roles in C_4_ biochemical carbon-concentrating mechanism were identified in sequenced Poaceae genomes with emphasis on the *Setaria*. Carbonic anhydrase (CaH) converts CO_2_ to bicarbonate (HCO_3_^−^) which is then pre-assimilated by phospho*enol*pyruvate carboxylase (PEPC) to form OAA in the mesophyll (Furbank and Taylor 1995; Schuler et al. 2016). The conversion of OAA to malate is catalyzed by malate dehydrogenase (MDH) and following the diffusion of malate through plasmodesmata to the bundle sheath, decarboxylation of malate is catalyzed by one of three different decarboxylating enzymes to form pyruvate and CO_2_ (Hatch and Burnell 1990). Finally, PEP is regenerated in the mesophyll by pyruvate orthophosphate dikinase (PPDK) (Aubry et al. 2011). Based on the decarboxylating enzymes, three biochemical subtypes of C_4_ photosynthesis were defined. These include NADP-dependent malic enzyme (NADP-ME), NAD-dependent ME (NAD-ME) and phospho*enol*pyruvate carboxykinase (PEPCK) (Gutierrez et al. 1974). The present study involves two subtypes, namely NADP-ME (*S. italica, S. viridis, Zea mays* and *Sorghum bicolor*) and NAD-ME (*Panicum hallii* and *P. virgatum*). Also, *Oryza sativa* and *Brachypodium distachyon* from C_3_ clade have been included. Since no genome of a PEPCK subtype has been sequenced, it has been excluded from the present study.

All five gene family members were identified using computational approaches in these eight species. However, importance has been given to analyze the genic, genomic and physicochemical protein properties of these gene family members in *S. italica* and *S. viridis*. The different classes of enzymes were further categorized according to their phylogeny, domain architecture, subcellular localization and sequence conservation. Promoter analysis has been performed to identify the *cis-*regulatory elements present in the upstream regions responsible for the regulation of gene expression at the transcriptional level. The RNA-seq data of different tissues and stress treatment were systematically analyzed to derive the expression profiles of these five gene family members. Based on this, candidate genes were chosen for expression profiling in response to different abiotic stresses (dehydration, salt and heat) and hormone treatments (abscisic acid, jasmonic acid and salicylic acid) in two contrasting *S. italica* cultivars. Comparative mapping of these five gene families between the sequenced grass genomes and Ks dating of orthologous gene pairs were performed to deduce the evolutionary aspects of these gene families.

## Materials and methods

### Identification of C_4_ photosynthetic genes

Three different strategies, namely, keyword, BLASTp and HMM (Hidden Markov Model) searches, were deployed to identify the C_4_ photosynthetic genes in sequenced Poaceae species. The names of proteins were used as a keyword to search the annotated protein database of Phytozome v12 (https://phytozome.jgi.doe.gov). The protein sequences of CaH, PEPC, PPDK, NADP-MDH and NADP-ME reported in the literature (Wang et al. 2009, 2016; Imran et al. 2016; Tao et al. 2016) were retrieved and BLASTp searched against the Poaceae database in Phytozome v12. The resultant protein sequences of all the four families and their corresponding query sequences were used to prepare HMM profile for each family. The profiles were BLASTp searched against the protein database of Poaceae species using HMMER v3.2.1(http://hmmer.org/) with an e-value cut-off of 1.0. The obtained protein sequences of CaH, PEPC, PPDK, NADP-MDH and NADP-ME families were validated by SMART (http://smart.embl-heidelberg.de/) and PFAM (https://pfam.xfam.org/) databases to confirm the presence of signature domains conserved to each family. The sequence information (gene, transcript and CDS), chromosomal position and gene orientation were retrieved for each protein from the. gff file of *S. italica* v2.2 and *S. viridis* v2.1 available in Phytozome v12. The genes identified in *S. italica* and *S. viridis* were prefixed with „*Si*‟ and „*Sv*‟, respectively, in further analyses.

### Chromosomal localization, gene structure and phylogenetic analysis

MapChart v2.32 was used to construct the physical map showing the chromosomal localization of identified genes (https://www.wur.nl/en/show/Mapchart.htm). The structure of each gene was predicted using GSDS server v2.0 (http://gsds.cbi.pku.edu.cn/index.php) by aligning the CDS sequence on the corresponding gene sequence to identify the positions of introns, exons and UTRs. The phylogenetic tree was constructed using MEGA X v10.0.1 (https://www.megasoftware.net/) using Neighbor-Joining method with 1000 bootstrap iterations. The protein sequences were further scanned using NCBI CDD search tool (https://www.ncbi.nlm.nih.gov/Structure/bwrpsb/bwrpsb.cgi) to study their domain architecture. The physicochemical properties of each protein were retrieved from the ProtParam tool of ExPASy server (https://web.expasy.org/protparam/). Subcellular localization of identified proteins was predicted using WoLF PSORT (http://www.genscript.com/wolf-psort.html), TargetP (www.cbs.dtu.dk/services/TargetP) and Predotar (https://urgi.versailles.inra.fr/predotar).

### Promoter analysis and *in silico* expression profiling

The upstream sequence (~ 2 kb) of each gene (from TSS) were retrieved from Phytozome and searched for *cis-*regulatory elements using the *cis-*Map feature of PlantRegMap database (http://plantregmap.cbi.pku.edu.cn/cis-map.php). The publicly available transcriptome data of *S. italica* namely, root (SRX128223), stem (SRX128225), leaf (SRX128224) and spica (SRX128226), and a drought stress library (SRR629694 – stress; SRR629695 – control) were retrieved from European Nucleotide Archive (http://www.ebi.ac.uk/ena). The reads were processed to generate RPKM values following Muthamilarasan et al. (2015), and the heat map was visualized using MeV v4.9 (http://mev.tm4.org/).

### Synteny and duplication analysis of C_4_ photosynthetic genes

The physical maps were manually inspected for tandem duplications, whereas segmental duplications were predicted using MCScanX toolkit (http://chibba.pgml.uga.edu/mcscan2/) with default parameters. The identified protein and gene sequences were BLAST searched against the database of sequenced grass species namely, sorghum, maize, rice, Brachypodium, wheat and switchgrass available in Phytozome. Sequences showing more than 80 % similarity (e-value<1e-05) were considered as orthologous gene pairs. Reciprocal BLAST was performed with default parameters to confirm the findings, and comparative maps were constructed using Circos v0.96 (http://circos.ca/). The synonymous (Ks) and non-synonymous (Ka) substitution rates of paralogous as well as orthologous gene pairs were estimated using CODEML program in PAML tool of PAL2NAL (http://www.bork.embl.de/pal2nal/). Further, time of duplication and divergence was determined using a synonymous mutation rate of λ substitutions per synonymous site per year as T = Ks/2λ (where λ = 6.5×10^−9^ for monocots; Lynch and Conery 2000).

### Expression profiling by qRT-PCR in response to stress and hormone treatments

Seeds of *S. italica* cultivars contrastingly tolerant to drought and salt stresses (cv. IC-4, tolerant; cv. IC-41, susceptible) were allowed to germinate under controlled conditions as described in Singh et al. (2016). Twenty-one-day-old seedlings were exposed to 250 mM NaCl (salt), 20% PEG 6000 (dehydration), 45°C (heat) and 100 μM each of abscisic acid (ABA), methyl jasmonate (MJ) and salicylic acid (SA), and whole seedlings were collected at 0 (control), 6 (early) and 12 (late) hours post-treatment. Total RNA was isolated from each sample using Trizol reagent, and the quality and quantity were ascertained following Singh et al. (2016). cDNA synthesis was performed using Thermo Scientific Verso cDNA synthesis kit following manufacturer‟s instructions and qRT-PCR was performed in StepOne™ Real-Time PCR Systems (Applied Biosystems, USA) as described in Singh et al. (2016) (Supplementary Table S1). Three technical replicates for each biological duplicate was maintained, and the transcript abundance normalized to the internal control *Act2* was analyzed using 2^−ΔΔCt^ method (Kumar et al. 2013). The PCR efficiency was calculated as by the default software itself (Applied Biosystems, USA).

## Results

### The C_4_ photosynthetic enzyme repertoire of Poaceae genomes

The C_4_ photosynthetic enzyme repertoire of sequenced grass species was identified using computational approaches. The number of CaH proteins ranged from 5 to 15 in C_4_ clade, and only 3 and 2 proteins were identified in *B. distachyon* and *O. sativa*, respectively (Fig. 1). In the case of PEPC and PPDK, the number was almost invariable throughout the species studied; however, *P. virgatum* and *Z. mays* contained double the amount of PEPC proteins (12 and 10, respectively). *B. distachyon* had a single PPDK protein, whereas *P. virgatum* possessed three proteins (Fig. 1). NADP-MDH proteins ranged from 10 to 18 in C_4_ clade, and in C_3_ clade, the number was less (5 and 9 in *B. distachyon* and *O. sativa*, respectively). The number of NADP-ME proteins was higher in *Z. mays* (11), followed by *S. italica* (9), and *S. bicolor* and *S. viridis* (8 each). Since, the focus of the present study was to understand the structure, organization, evolution and expression pattern of C_4_ photosynthetic enzymes in *S. italica* and *S. viridis*, the enzymes and their encoding genes were studied elaborately in these two crops. An almost similar number of proteins were identified in both the crops (Fig. 1; Supplementary Table S2). The recent divergence of *S. italica* and *S. viridis* (5900 – 8900 years ago) has resulted in higher collinearity between their genomes (>95 %; Zhang et al. 2012), and therefore, the properties of these gene families were identical (Supplementary Table S2). Also, the physical maps showing the location of all the five gene family members in *Setaria* genomes were almost similar with minor chromosomal rearrangements (Fig. 2).

**Fig. 1.**
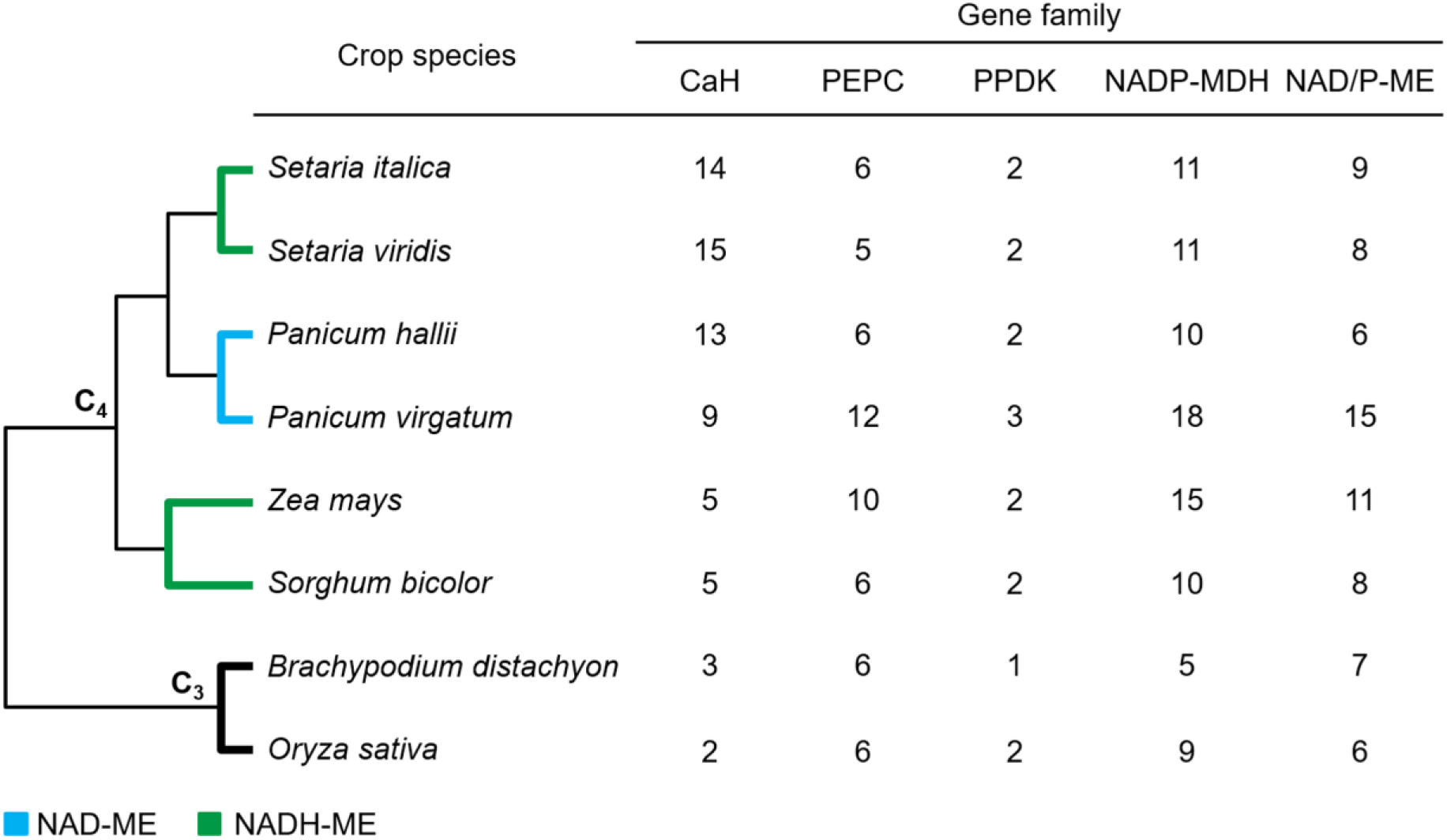
The distribution of C_4_ photosynthetic genes among sequenced Poaceae genomes. The relationship tree is showing the number of genes encoding carbonic anhydrase (CaH), phospho*enol*pyruvate carboxylase (PEPC), pyruvate orthophosphate dikinase (PPDK), NADP-dependent malate dehydrogenase (MDH) and NADP-dependent malic enzyme (NADP-ME) in the Poaceae genomes.

**Fig. 2.**
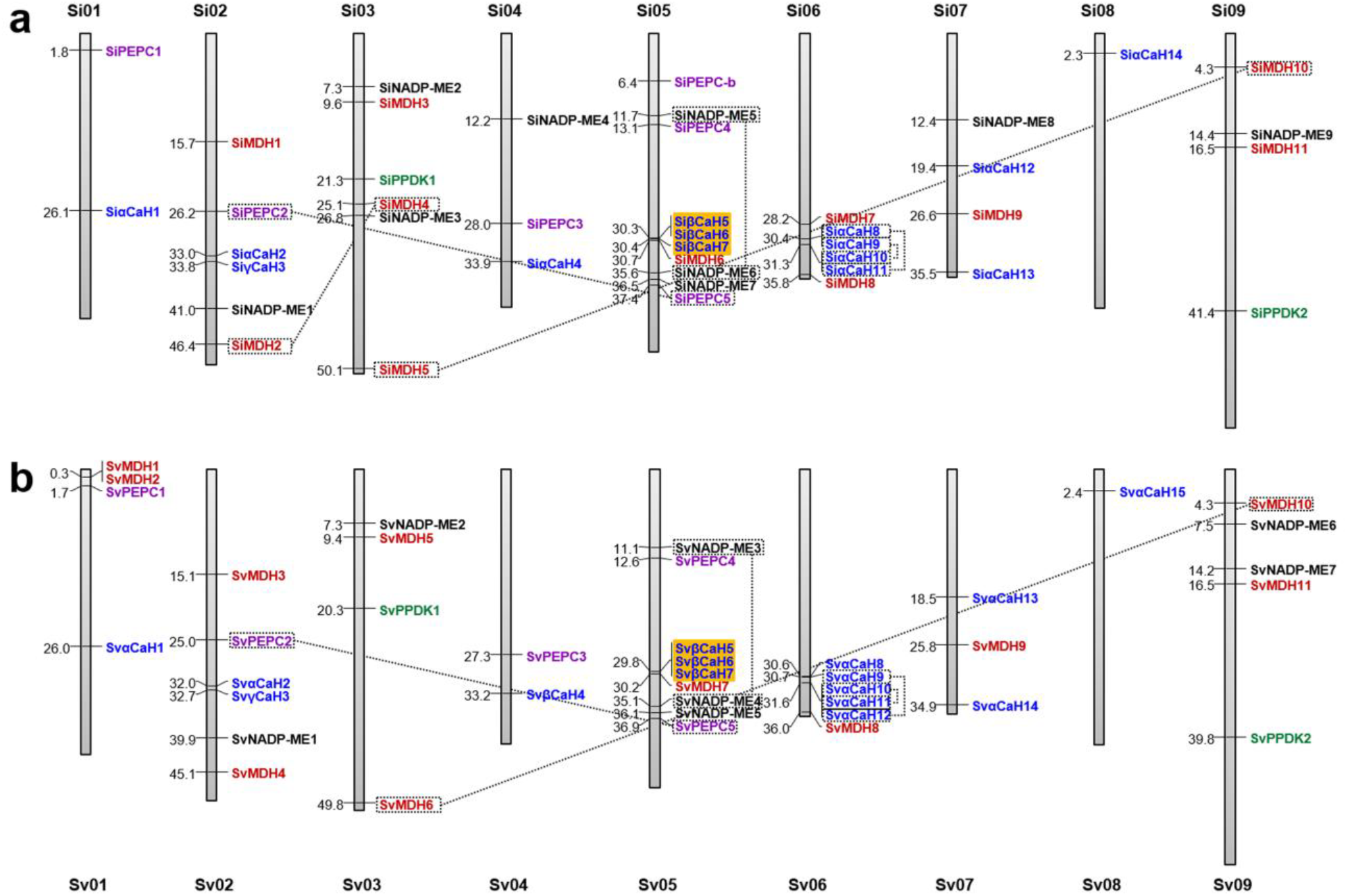
Chromosomal distribution of C_4_ photosynthetic genes in (**a**) *Setaria italica* and (**b**) *S. viridis*. Vertical bars represent chromosomes as labeled and the values on the left indicate the chromosomal location (in Mb). Segmentally duplicated genes are shown within dotted boxes, and dotted lines connect the gene-pairs. Tandemly duplicated genes are shown in the colored box.

### The carbonic anhydrase family of *Setaria*

A total of 14 and 15 CaH proteins were identified in *S. italica* and *S. viridis*, respectively. The proteins were smaller in size (~110 – 330 amino acids) with a molecular weight range of ~12 – 36 kDa. Phylogenetic analysis of SiCaH and SvCaH proteins showed three distinct subclasses, namely, α, β and γ (Fig. 3a). The α-CaH subclass contained a maximum of 10 and 11 proteins in *S. italica* and *S. viridis*, respectively; whereas, the β- and γ-CaH subclasses possess 1 and 3 proteins, respectively, in both the crops. The α- and γ-CaH proteins contained the „alpha_CA domain of prokaryotic origin‟, and in the case of β-CaH, „beta_CA domain‟ was present. Further, subcellular localization predicted the localization of α-CaH in the extracellular matrix, vacuole and chloroplast, whereas β-CaH proteins were present only in the chloroplast. On the contrary, γ-CaH was mitochondria-localized (Supplementary Table S2). The mitochondrial localization of γ-CaH could be attributed to its role in maintaining mitochondrial physiology, including the formation of complex I assembly (Meyer et al. 2011). Also, γ-CaH plays a role in male sterility (Villarreal et al. 2009), and growth and embryogenesis (Fromm et al. 2016). The α-CaH were reported to be involved in physiological processes in aboveground tissues (Villarejo 2005; Bhat et al. 2017). The β-CaH has a major role in C_4_ photosynthesis as it performs hydration of CO_2_ to form HCO_3_^−^ which acts as a substrate for PEPC to synthesize oxaloacetate (Ludwig 2016). Thus, PEPC along with CaH is imperative for establishing rapid equilibrium between CO_2_ and HCO_3_^−^ to facilitate carbon fixation.

**Fig. 3.**
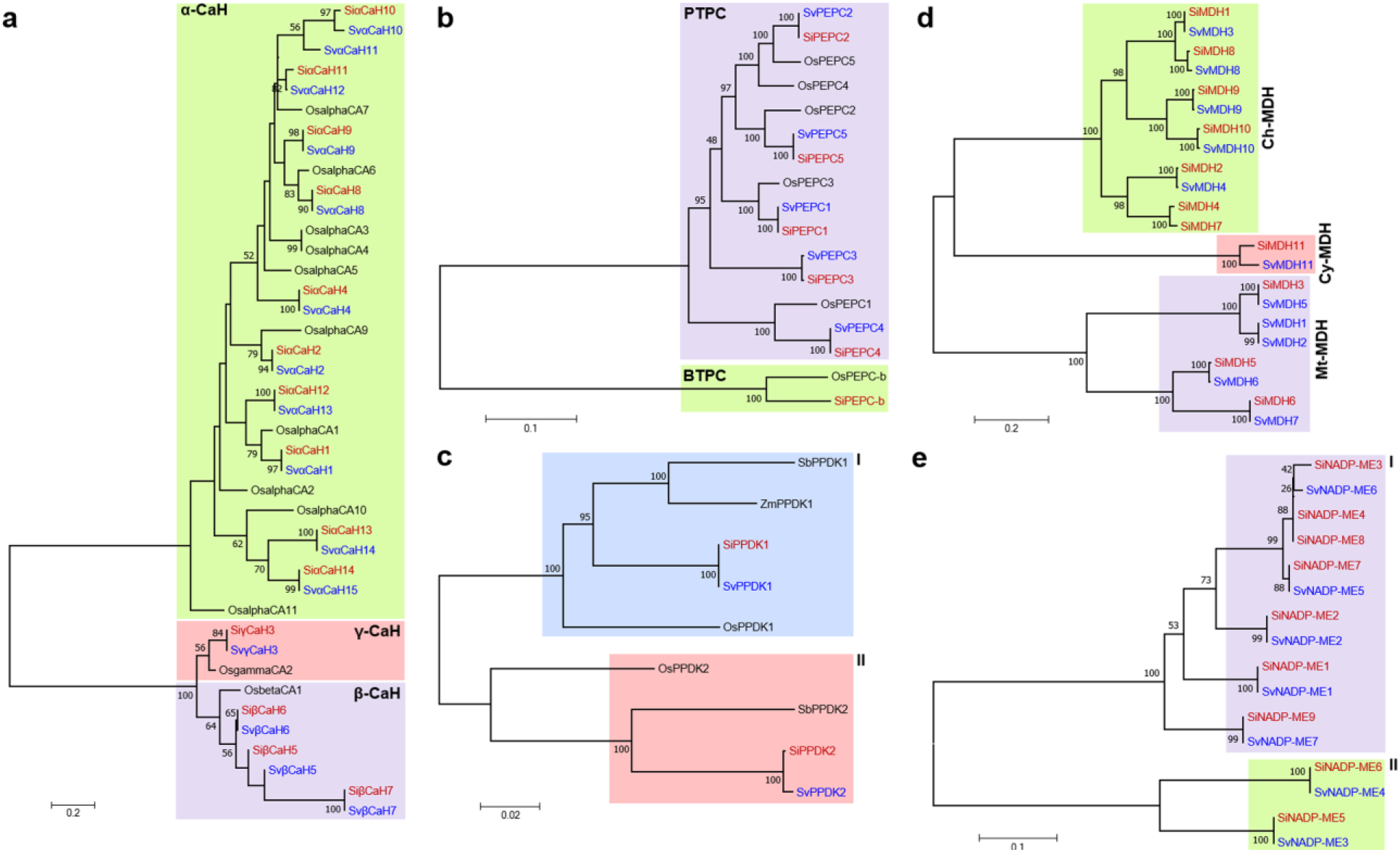
Phylogenetic relationships of C_4_ photosynthetic enzymes namely, (**a**) carbonic anhydrase, (**b**) phospho*enol*pyruvate carboxylase, (**c**) pyruvate orthophosphate dikinase, (**d**) NADP-dependent malate dehydrogenase and (**e**) NADP-dependent malic enzyme. The amino acid sequences of C_4_ photosynthetic enzymes of *S. italica* and *S. viridis* were aligned using ClustalW, and the alignment file was used to construct the Neighbor-Joining tree with 1000 bootstrap iterations under default parameters.

In terms of gene size and structure, the CaH genes were 0.5 to ~7 kb in length (Supplementary Table S2). The shortest genes, *SiαCaH13* and *SvαCaH14*, were intronless, while *SiβCaH7* and *SiγCaH3* of *S. italica* and *SvαCaH7* and *SvγCaH3* of *S. viridis* had a maximum of seven introns (Supplementary Figure S1). Chromosomal localization of *CaH* genes showed their distribution in all the chromosomes except chromosome 3 and 9 (Fig. 2). Maximum genes were found to be localized on chromosomes 5 and 6 of both the crops and interestingly, chromosome 5 possessed only the genes of βCaH subclass and chromosome 6 contained αCaH subclass. Among these, βCaH subclass members on chromosome 5 were found to be tandemly duplicated, whereas the αCaH subclass members on chromosome 6 were segmentally duplicated (Fig. 2). This shows that tandem and segmental duplications have contributed to the evolution of CaH gene family in *Setaria*. In sorghum, the tandem gene cluster of βCaH subclass was reported on chromosome 3 (Wang et al. 2009), which shows that the duplication has occurred before the divergence of these species.

### The phospho*enol*pyruvate carboxylase family of *Setaria*

In the C_4_ photosynthetic pathway, PEPC catalyzes the irreversible β-carboxylation of phospho*enol*pyruvate in the presence of HCO_3_^−^ to produce oxaloacetate and inorganic phosphate (Izui et al. 2004). A total of 6 and 5 PEPC proteins were identified in *S. italica* and *S. viridis*, respectively (Supplementary Table S2). *S. italica* possessed an additional member (SiPEPC-b) belonging to bacterial-type PEPC (BTPC). The other five PEPCs of both the crops belong to plant-type PEPC (PTPC). The same was reflected in the phylogenetic tree, where PTPCs formed a separate clade, and BTPC of rice and *S. italica* formed a distinct clade (Fig. 3b). Analysis of domain architecture revealed that SiPEPC-b contained a prokaryotic-like (R/K) NTG C-terminal tetrapeptide (PRK00009) domain (Supplementary Table S3). Interestingly, this protein was larger (1032 amino acids; 115 kDa) and exhibited less sequence identity with PTPCs (<50%). Similar observations were reported in soybean (Wang et al. 2016), Arabidopsis (Sánchez and Cejudo 2003), rice (Sánchez and Cejudo 2003), lotus (Nakagawa et al. 2003) and wheat (Duff and Chollet 1995); however, no report is available on BTPC and PTPC in C_4_ grasses. O’Leary et al. (2011a,b) have shown that BTPCs occur as catalytic and regulatory subunits of extraordinary heteromeric complexes (Class-2 PEPCs).

The PTPCs of *S. italica* and *S. viridis* had a close size range of 964 (~110 kDa) to 1015 amino acids (~114 kDa) with a conserved N-terminal seryl-phosphorylation site, and these proteins could characteristically exist as homotetrameric Class-1 PEPCs (O’Leary et al. 2011b). The sizes of PEPC encoding genes ranged from ~5 to 9 kb in both the crops with a distribution of introns ranging from 6 - 9 (Supplementary Figure S1). With an exception, SiPEPC-b had 19 introns which agree to BTPCs reported in other crops including sorghum (Wang et al. 2009), soybean (Wang et al. 2016) and rice (Sánchez and Cejudo 2003). Chromosomal localization of PEPC genes showed their localization on chromosomes 1, 2, 4 and 5, where the genes *PEPC2* (chromosome 2) and *PEPC5* (chromosome 5) were found to be segmentally duplicated in both the crops (Fig. 2).

### The pyruvate orthophosphate dikinase family of *Setaria*

PPDK catalyzes a reversible conversion of pyruvate to phospho*enol*pyruvate in an ATP- and Pi-dependent manner (Aubry et al. 2011). Two proteins were identified in *S. italica* and *S. viridis*, which is in congruence to the numbers reported in sorghum, maize and rice (Wang et al. 2011). These two proteins, designated as PPDK1 and PPDK2, form two distinct clades in the phylogenetic tree (Fig. 3c). In terms of gene length and structure, *PPDK1* were ~14 kb in size with 18 introns, and *PPDK2* were ~8 kb with 16 (*S. italica*) and 17 (*S. viridis*) introns (Supplementary Table S2; Supplementary Figure S1). Physical map showed the localization of *PPDK1* and *PPDK2* on chromosomes 3 and 9, respectively (Fig. 2). Though length, molecular weight and domain architecture of these proteins were almost similar, all the PPDKs were predicted to be localized in chloroplast except SiPPDK2 which was cytoplasm-localized (Supplementary Table S2; Supplementary Table S3). However, the presence of chloroplast transit peptide (cTP) of 45 amino acids length was observed only in SiPPDK1 and SvPPDK1 that plays a role in transporting these proteins to chloroplast stroma (Soll and Tien 1998). Thus, further experimental validations are required to confirm the subcellular localization of these proteins.

### The NADP-dependent malate dehydrogenases of *Setaria*

In C_4_ crops, MDH mediates the interconversion of oxaloacetate to malate by utilizing the NAD/NADH coenzyme system (Hatch and Burnell 1990; Furbank and Taylor 1995; Schuler et al. 2016). Eleven MDH proteins were identified in both *S. italica* and *S. viridis* with similar structural properties (Supplementary Tables S2). The size of the proteins ranged from 332 amino acids (~35 kDa) to 452 amino acids (~50 kDa), and their isoelectric pH ranged from 5.75 to 8.71. Phylogenetic analysis classified these proteins into three distinct subgroups according to their subcellular localization properties, namely chloroplast-MDH (Ch-MDH), cytoplasmic-MDH (Cy-MDH) and mitochondrial-MDH (MtMDH) (Fig. 3d). The analysis revealed that predominant proteins of *S. italica* were chloroplast localized (7) followed by mitochondria (3); however, in *S. viridis*, an equal number of proteins were found to be localized in both the organelles (5 each). One protein each of *S. italica* and *S. viridis* was found to be localized in the cytoplasm. Similar classification based on sequence similarities and sequences conserved for subcellular localization has been reported in *G*. *raimondii*, *G arboretum* and *G*. *hirsutum* (Imran et al. 2016).

The sizes of *MDH* genes ranged from ~1 kb to ~5 kb in both the crops; however, their structure and chromosomal distributions were different, unlike other gene families (Supplementary Table S2). In *S. viridis*, the two genes were uniquely mapped on chromosome 1 beside their localization on chromosomes 2, 3, 5, 6, 7 and 9 in both the crops (Fig. 2). In *S. italica*, *SiMDH2* (chromosome 2) and *SiMDH4* (chromosome 3), and *SiMDH5* (chromosome 3) and *SiMDH10* (chromosome 9) were segmentally duplicated; whereas only the latter was found in *S. viridis*. Two genes of *S. italica* (*SiMDH2* and *SiMDH4*) were intronless, but in case of *S. viridis*, only *SvMDH2* was intronless (Supplementary Figure S1). The highest number of introns were found to be present in *SiMDH1, SiMDH8, SvMDH2* and *SvMDH8* genes (13 introns).

### The NADP-dependent malic enzymes of *Setaria*

The NADP-dependent malic enzyme catalyzes the formation of pyruvate from malate with the release of CO_2_ (Chang and Tong 2003). Nine and eleven NADP-ME proteins were identified in *S. italica* and *S. viridis*, respectively. Unlike other families, NADP-ME possessed a broad range of size and molecular weight. The lengths of these proteins ranged from 149 to 652 amino acids with a molecular weight of 16 to 72 kDa (Supplementary Tables S2). Similarly, the pI ranged from 5.4 to 8 in both the crops. Phylogenetic classification revealed the two distinct subgroups (I and II) among the NADP-ME gene family (Fig. 3e). Phylogenetic classification of this gene family based on subcellular localization was previously reported in dicots (Wheeler et al. 2005); however, no such distinct categorization has been formulated so far in monocots (Chi et al. 2004; Rao and Dixon 2016). In the present study, the members of the subgroup I were predicted to be localized in mitochondria, whereas the members of subgroup II were chloroplast localized (Fig. 3e). The members of subgroup II might have roles in C_4_ photosynthetic pathway, particularly in the bundle sheath chloroplasts where they perform the oxidative decarboxylation of malate to produce pyruvate, CO_2_, and NADPH in the presence of a divalent cation (Chang and Tong 2003; Rao and Dixon 2016).

Similar to MDH genes, the members of NADP-ME gene family were also relatively divergent between *S. italica* and *S. viridis* in terms of genomic properties (Supplementary Table S2). The size of NADP-ME genes ranged from ~2 to 8 kb in *S. italica* and ~1 to 7 kb in *S. viridis.* The physical map showed their localization on all the chromosomes except 1, 6 and 8 in *S. italica* (Fig. 2). In the case of *S. viridis*, the genes were localized only on chromosomes 2, 3, 5 and 9. Segmental duplication of two genes on chromosome 5 in *S. italica* (*SiNADP-ME5* and *SiNADP-ME6*) and *S. viridis* (*SvNADP-ME3* and *SvNADP-ME4*) was also observed (Fig. 2).

### *Cis-*elements in the promoter regions of C_4_ photosynthetic genes

A total of 250 *cis-*regulatory elements were present on the 2 kb upstream of the five gene families in both the crops (Supplementary Table S4). Similar to sequence conservation between the genic regions of *S. italica* and *S. viridis*, the upstream regions were also found to highly conserved, and therefore, there was no much difference between the elements identified for the same gene in both the crops. In this study, several vital motifs were identified including; (i) I-box (IBOXCORE), a conserved motif present in the upstream of light-regulated genes; (ii) „CBFHV‟, site for dehydration-responsive element (DRE) binding protein (DREB); (iii) GT-1 motif (GT1GMSCAM4), which regulates pathogen- and salt-induced *SCaM-4* gene expression; (iv) Core of low temperature responsive element (LTRECOREATCOR15); (v) „MYB1AT‟ and „MYCCONSENSUSAT‟, MYB recognition sites present in dehydration-responsive genes; and (iv) GATA box, a motif required for advanced tissue-specific expression regulated by light. In addition, several hormone-regulated motifs were also identified including (i) „ABRELATERD1‟, an abscisic acid responsive element-like sequence required for early response to dehydration; (ii) „RAV1AAT‟, an ethylene-responsive element; (iii) GCC-box (GCCCORE) associated with ethylene-regulated dehydration response; (iv) „ARR1AT‟, an auxin-responsive element; (v) „GAREAT‟, a gibberellic acid responsive element; (vi) CGTCA- and TGACG-motifs for jasmonic acid response; and (vii) TCA-element, a salicylic acid regulatory motif.

In addition, there were several elements unique to one or two members of C_4_ photosynthetic families. This includes (i) ACIII element (ACIIIPVPAL2) present only in *NADP-ME6*, which is required for vascular-specific gene expression; (ii) cell cycle box (CELLCYCLESC), responsible for cell-cycle-specific activation of transcription; (iii) E2Fa element (E2FANTRNR), the binding site of E2F present in *βCaH7* and *PEPC5*; (iv) M-specific activator (MSACRCYM) in *αCaH13* essential for M phase-specific expression; (v) Palindromic C-box (PALINDROMICCBOXGM) for bZIP transcription factor binding; and (vi) Wound-responsive element (WRECSAA01) (Supplementary Table S4). The presence of diverse elements at the upstream region of C_4_ photosynthetic genes suggests the involvement of these genes in several molecular and physiological functions. However, further experiments are necessary to validate these findings and pinpoint the precise elements that have roles in the trait of interest.

### RNA-Seq derived expression profiles of C_4_ photosynthetic genes in tissues and dehydration stress

The RNA-seq data of four tissues of *S. italica* namely root, leaf, spica and stem as well as the dehydration library were analyzed to derive the expression values (RPKM) for each gene, and the heatmap was generated (Fig. 4a). Overall, the heatmap showed a differential expression pattern in all the genes with few distinct expression profiles. The data showed tissue-specific upregulation in few genes, including *SiMDH6* (spica and root), *SiMDH10* (leaf) and *SiNADP-ME6* (root). Several genes including *SiβCaH5*, *SiPPDK1*, *SiMDH8* and *SiNADP*-*ME5* were upregulated in all the tissues except root. Interestingly, *SiPPDK2* did not show any expression in all the libraries. *SiMDH3* showed higher expression in all the tissues except leaf. Comparing the tissue-specific expression data with stress library highlighted the putative stress-responsive genes (Fig. 4a). *SiγCaH3*, *SiPEPC1*, *SiPEPC2* and *SiMDH5* showed a distinct upregulation during dehydration, whereas *SiMDH2* and *SiNADP-ME4* were downregulated in response to stress treatment. There were few genes which did not show any expression in both the tissues as well as stress sample, and this includes *SiαCaH1, SiαCaH14, SiPEPC-b, SiMDH4, SiMDH9, SiNADP-ME3* and *SiNADP-ME8*.

**Fig. 4.**
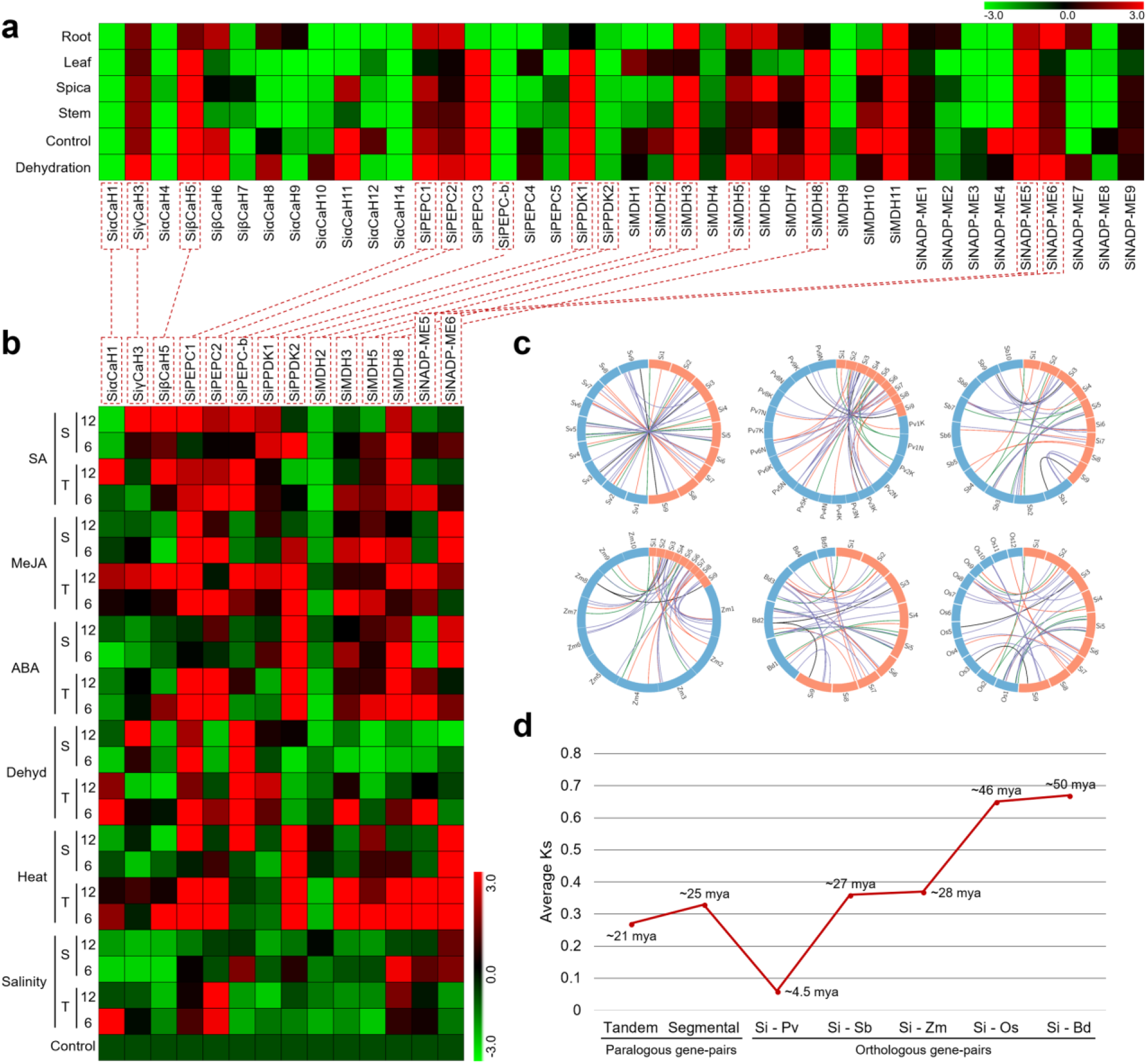
Expression analysis and comparative mapping of C_4_ photosynthetic genes. (**a**) Heatmap showing the expression pattern of C_4_ photosynthetic genes in four tissues and dehydration library. (**b**) Relative expression of candidate C_4_ photosynthetic genes in response to different abiotic stresses (dehydration, salt and heat) and hormone treatments (abscisic acid, jasmonic acid and salicylic acid) at three timepoints (0 h, 6 h and 24 h) in two contrasting *S. italica* cultivars [„T‟ and „S‟ indicate the tolerant (IC-4) and susceptible cultivars (IC-41), respectively]. The heatmap is generated based on the fold-change values in the treated sample compared to control (0 h) sample. The color scale for fold-change values is shown at the top, where −3.0, 0.0 and 3.0 denote low, medium and high expression, respectively. (**c**) Comparative physical mapping of orthologous gene-pairs between *S. italica* (Si) and *S. viridis* (Sv)*, P. virgatum* (Pv), *S. bicolor* (Sb), *Z. mays* (Zm), *B. distachyon* (Bd), and *O. Sativa* (Os). Circular blocks represent chromosomes, and colored lines connect the orthologous gene-pairs. (**d**) Time of duplication and divergence (mya) estimated based on synonymous substitution rate (Ks) of paralogous and orthologous gene-pairs.

These genes may have a role in growth, development or response to other biotic and abiotic stresses, which has to be studied further. Based on the RNA-Seq derived expression profile, fourteen genes were chosen for analyzing their expression in response to abiotic stresses and hormone treatments in *S. italica*.

### Expression profiles of C_4_ photosynthetic genes in stress and hormone treatments

The transcript abundance of fourteen candidate C_4_ photosynthetic genes was analyzed using qRT-PCR in two contrasting cultivars of *S. italica* at early and late time-points of stress and hormone treatments (Fig. 4b). During salinity stress, relatively no difference was observed in the expression profiles of all the genes except the upregulation of *SiαCaH1* at the early stage of stress in the tolerant cultivar (IC-4), *SiPEPC2* at both the time-points in the tolerant cultivar and *SiMDH8* at the early stage of susceptible cultivar (IC-41). In case of heat stress, the genes *SiβCaH5* and *SiMDH5* showed higher expression at the early stage of stress treatment in the tolerant cultivar, whereas *SiPEPC2*, *SiMDH3*, *SiMDH8* and *SiNADP*-*ME5* were upregulated at both the timepoints in this cultivar. On the contrary, *SiPEPC1* and *SiPEPC*-*b* showed higher expression at 12 h post-stress treatment in the susceptible cultivar.

The genes, *SiPPDK2* and *SiNADP-ME6* were highly expressed in all the time-points in both the cultivars. Dehydration stress did not significantly alter the transcript level of C_4_ photosynthetic genes; however, *SiPEPC*-*b* showed higher expression in all the time-points in both the cultivars. Upregulation in the expression of *SiPEPC1* at the early time-point of stress treatment in the tolerant cultivar was also observed. Also, *SiαCaH1*, *SiMDH3* and *SiNADP-ME5* were found to be highly expressed at the early stage of stress treatment in the tolerant cultivar. Only *SiγCaH3* showed upregulation in the susceptible cultivar at 12 h post-dehydration stress.

Similar expression patterns were detected among these genes during hormone treatments where differential expression of several genes suggests their regulation by these hormones in modulating cellular and molecular activities *in planta* (Fig. 4b). Among CaH gene family members, *SiαCaH1* did not show a notable change in its expression levels during hormone treatments except at 12 h of SA in the tolerant cultivar. In contrast, *SiγCaH3* and *SiβCaH5* were upregulated at 12 h post-MeJA and SA treatments in the tolerant and susceptible cultivars, respectively. Higher expression of PEPC gene family members was observed during hormone treatments. *SiPEPC1* and *SiPEPC2* showed distinct upregulation during ABA treatment in the tolerant cultivar, whereas *SiPEPC-b* was found to be modulated only during SA treatment in both the cultivars. *SiPPDK2*, in contrast to *SiPPDK1*, showed higher expression during ABA (in all the time-points of both the cultivars), MeJA (in all the time-points of tolerant cultivar) and SA (6 h post-SA treatment in the susceptible cultivar). The expression of *SiMDH2* was unaffected by hormones; however, *SiMDH3* and *SiMDH5* showed an insignificant change in their expression in response to all the stresses in both IC-4 and IC-41. In the case of NADP-MEs, *SiNADP-ME5* was upregulated during early time-points of ABA and SA, and late timepoint of MeJA in the tolerant cultivar and its expression remained unaltered in the susceptible cultivar. Contrarily, *SiNADP-ME6* showed higher expression in both the timepoints of ABA and MeJA treatments in the susceptible cultivar.

### Divergence of C_4_ photosynthetic genes among Poaceae members

To gain understanding on the organization and evolution of C_4_ photosynthetic genes among sequenced Poaceae genomes, orthologous genes of *S. italica* were identified in *P. virgatum, S. bicolor, Z. mays, O. sativa* and *B. distachyon* (Fig. 4c). Of the 42 genes of *S. italica*, 28 genes were orthologous to *P. virgatum* (~67 %), 25 to *S. bicolor* (~60 %), 26 to *Z. mays* (~62 %), 18 to *O. sativa* (~43 %) and 15 to *B. distachyon* (~36 %) (Supplementary Table S5). Among CaH gene family, *SiαCaH1* was conserved in all the grass species studied, whereas orthologs of *SiαCaH2* and *SiαCaH4* were present in all the crops except rice. Comparatively, 9 genes of CaH gene family were orthologous to *P. virgatum*, followed by *S. bicolor* and *Z. mays* (5 each), *B. distachyon* (4 genes) and *O. sativa* (2 genes). In case of PEPC gene family, *SiPEPC1* to *SiPEPC5* were conserved among *S. bicolor* and *Z. mays*, whereas only four genes were orthologous to *O. sativa*, three to *P. virgatum* and two to *B. distachyon*. PPDK family members were found to be highly conserved throughout the grass genomes as the orthologs were identified in all the five species studied. Similarly, nine MDH genes showed an orthologous relationship with *S. bicolor*, eight with *P. virgatum* and *Z. mays*, five and six genes with *O. sativa* and *B. distachyon*, respectively. Among these, *SiMDH1*, *SiMDH3* and *SiMDH9* were conserved among all the five genomes. The members of NADP-ME gene family showed maximum synteny with *P. virgatum* and *Z. mays* (6 gene pairs each), followed by *S. bicolor* and *O. sativa* (4 gene pairs each), and *B. distachyon* (3 gene pairs) (Supplementary Table S5).

To explore the selective constraints on duplication and divergence of C_4_ photosynthetic genes, the ratios of non-synonymous (Ka) versus synonymous (Ks) substitution rate (Ka/Ks) for paralogous and orthologous gene-pairs were determined (Fig. 4d; Supplementary Table S5). In the case of duplicated gene pairs, the average Ks values were 0.07 and 0.13 for *S. italica* and *S. viridis*, respectively, which indicated that these genes underwent intense purifying selection pressure. Further, the estimation of their time of duplication revealed that the tandem and segmental duplications occurred around 21 and 25 million years ago (mya). This is in agreement with the whole genome duplications reported in *S. italica* and *S. viridis* predicted using genome sequence data (Zhang et al. 2012). In case of orthologous gene pairs, the average Ks values were 0.06, 0.36, 0.37, 0.65 and 0.67 for *S. italica – P. virgatum, S. italica – S. bicolor, S. italica – Z. mays, S. italica – O. sativa* and *S. italica – B. distachyon*, respectively (Fig. 4d). This showed that these species underwent strong purifying selective pressure. Further, the data also showed that *S. italica – P. virgatum* divergence has recently occurred, around 4.5 mya, and this divergence has happened after the whole genome duplication events as suggested by the data of paralogous gene-pairs. The divergence of *S. italica – S. bicolor* and *Z. mays* were predicted to be around 27-28 mya, whereas *S. italica – O. sativa* and *S. italica – B. distachyon* divergence has occurred about 46 and 50 mya, respectively. The data indicate that the C_4_ photosynthetic genes had undergone several duplication and divergence events, which might have enabled them to have different functional roles in physiological, molecular and stress-responsive machinery.

## Discussion

In the present study, a higher number of CaH genes was observed in *S. viridis* (15), *S. italica* (14) and *P. virgatum* (13) and the lowest were in C_3_ species including *B. distachyon* (3) and *O. sativa* (2) (Fig. 1). Among these, C_4_ species contained β-CaH subclass that was chloroplast localized. This suggests the involvement of these proteins in the conversion of CO_2_ to HCO_3_^−^ in the chloroplast of mesophyll cells. Besides, the roles of CaH in other physiological and biochemical processes have been reported in both C_3_ and C_4_ species. In *Arabidopsis*, *Atβ*CA1 and *Atβ*CA4 were reported to play roles in stomatal development and movement (Furbank and Taylor 1995; Schuler et al. 2016). The involvement of CaH proteins in nodule development and function was observed in alfalfa (Coba de la Peña et al. 1997), soybean (Kavroulakis et al. 2000) and lotus (Flemetakis et al. 2003). Expression profiling of *SiCaH* genes in response to stress treatments at early and late time-points of two different cultivars of *S. italica* showed their differential expression pattern (Fig. 4b). Significant upregulation of *SiαCaH1* was observed in the tolerant cultivar during salt and dehydration stress. Similarly, maize CaH gene, *NM_001111889* showed higher expression in salinity-induced cDNA subtraction library (Kravchik and Bernstein 2013). In rice, *OsCA1* showed at least two-fold upregulation in response to salt and osmotic stress and overexpressing this gene in *Arabidopsis* conferred tolerance to salinity stress (Yu et al. 2007). Unlike other *CaH* genes tested, *SiβCaH5* showed higher expression during heat stress alone (45°C) in the tolerant cultivar, which suggests its putative role in molecular response to high temperatures. However, no report so far indicates the involvement of *CaH* genes in heat and further experimentations are required to elucidate their precise role in heat stress tolerance.

The number of *PEPC* genes were almost the same in all the species except *P. virgatum* (12) and *Z. mays* (10). Irrespective of the number, one or two genes were reported to function in C_4_ photosynthesis so far. Other non-photosynthetic roles of PEPC include seed development and germination (O’Leary et al. 2011a), fruit ripening (O’Leary et al. 2011b) and photosynthetic isotope exchange and stomatal conductance (Cousins et al. 2007). Further, Christin et al. (2013) have amplified the PEPC genes of *O. sativa*, *S. bicolor*, *Z. mays* and few other representative species from PACCAD clade and shown that *ppc-C_4_* and *ppc-B2* were involved in C_4_ photosynthesis. Orthologs of these genes that have demonstrated differential expression in RNA-Seq derived expression data were chosen for qPCR analysis to deduce their transcript abundance during abiotic stresses. The results showed a higher expression pattern of these genes in response to salinity, heat and dehydration stresses specifically in the tolerant cultivar of *S. italica* (Fig. 4b). Similar upregulated expression of *PEPC* genes in response to salinity and/or drought stress has been reported in sorghum (García-Mauriño et al. 2003), wheat (González et al. 2003), lotus (Nakagawa et al. 2003) and tobacco (Doubnerová Hýsková et al. 2014). An increase in PEPC enzyme activity in response to drought stress was observed in three C_4_ subtypes namely, *Paspalum dilatatum* (NADP-ME), *Cynodon dactylon* (NAD-ME) and *Zoysia japonica* (PEPCK) suggesting their role in increased CO_2_ assimilation during water deficit conditions (Chang et al. 2012).

In C_4_ photosynthesis, PPDK performs the primary role of PEP and CO_2_ regeneration. PPDK gene family is the smallest among the other, with one to two members (Fig. 1). One of the two genes possesses two functionally independent promoters for generating a larger transcript with cTP expressed in leaves and a smaller transcript without cTP expressed in reproductive organs (Furbank and Taylor 1995; Wang et al. 2009; Schuler et al. 2016). The same has been observed in the present study, which suggests that the dual-promoter system is conserved in C_3_ and C_4_ species. In RNA-seq derived expression profile, *SiPPDK1* showed high expression in leaf, stem and spica, as well as control and dehydration stressed samples, whereas *SiPPDK2* did not show any difference in its expression in all the libraries (Fig. 4a). Similarly, qPCR data indicated that *SiPPDK1* did not show any notable difference in its expression during salt and heat stress in both tolerant and susceptible cultivars. However, *SiPPDK2* was highly expressed in both the cultivars exposed to heat stress (Fig. 4b). Transcriptome sequencing of mesophyll and bundle sheath cells of maize (Chang et al. 2012) and sorghum (Döring et al. 2016) has shown higher transcript abundance of *PPDK* in mesophyll cells. In another study, overexpression of maize *PPDK* has been shown to enhance the photosynthetic efficiency in wheat (Zhang et al. 2014). These evidence suggest the intriguing roles of PPDK in improving photosynthetic characteristics of C_4_ as well as C_3_ plants.

The activity of NADP-MDH depends on its subcellular localization. The leaf respiration and photorespiration were controlled by mitochondrial MDH (Tomaz et al. 2010), whereas fatty acid β-oxidation is regulated by peroxisomal MDH (Pracharoenwattana et al. 2007). The plastidic MDH plays two different roles in C_3_ and C_4_ plants. In C_3_ plants, MDH regulates the NADPH/ATP balance (Scheibe 1987), and in the C_4_ system, it catalyzes malate production (Berkemeyer et al. 1998). In the present study, expression levels of two plastidic (*SiMDH2* and *SiMDH8*) and mitochondrial MDHs (*SiMDH3* and *SiMDH5*) were analyzed in response to abiotic stresses. Interestingly, *SiMDH2* did not show a notable difference in its expression during stress as well as hormone treatments, while *SiMDH8* showed upregulation only during heat stress in the tolerant cultivar (Fig. 4b). Expression pattern of mitochondrial MDHs was also similar to plastidic MDHs, suggesting the involvement of these genes in heat stress response. Reports have shown the role of MDHs in improving cold tolerance in transgenic plants, for example, overexpression of apple cytosolic malate dehydrogenase gene in apple and tomato conferred tolerance to cold (Wang et al. 2016). Thus, it is suggested to analyze the transcript abundance of all the *SiMDH* genes in *S. italica* exposed to different abiotic stress treatments to identify potential candidates for overexpression studies.

NAD/P-ME catalyzes the decarboxylation of malate in the C4 cycle. In addition to its role in C_4_ photosynthesis, the enzyme is found to be involved in defense responses (Casati et al. 1999), lipid biosynthesis (Eastmond et al. 1997) and control of stomatal closure (Laporte et al. 2002). The expression levels of two *SiNADP-ME* genes were analyzed, which showed that both the genes are suggestively involved in heat stress response. However, *SiNADP-ME5* also showed a significant upregulation at the early stage of dehydration stress in the tolerant cultivar. Tao *et al*. (2016) have identified NADP-ME family members in 12 crucifer species and showed that the genes were involved in cold stress response. In rice, four genes were shown to be differentially expressed in response to salt and dehydration stresses (Chi et al. 2004). These observations demonstrate that NADP-ME family members could be putative candidates for further functional characterization to elucidate its molecular roles in C_4_ photosynthesis as well as stress tolerance.

## Conclusions

The C_4_ photosynthetic pathway has evolved from the C_3_ pathway to overcome the limitation of photorespiration by deploying a sophisticated biochemical carbon-concentrating mechanism. Keeping this in view, a comprehensive analysis has been performed to identify and characterize the pathway genes. Five key gene families were analyzed that provided insights into structure, organization, duplication and divergence, and their expression profiles in response to stress and hormone treatments. Differential expression pattern of a gene in contrasting cultivars suggest the presence of distinct regulatory mechanisms in the tolerant cultivar or sequence-level variations between the cultivars that should be studied further to understand the gene functional properties at the species level. The promoter analysis data defined in the present study will assist in executing this analysis. Comparative expression analysis of these genes in response to stresses in C_3_ and C_4_ crops followed by studying their localization and interaction with other proteins will advance our knowledge on understanding the fundamental difference between two pathways. The comparative maps constructed in the present study will aid the selection of candidate orthologous gene pairs. The genes identified in this study, namely, *SiαCaH1*, *SiβCaH5*, SiPEPC2, *SiPPDK2*, *SiMDH8* and *SiNADP-ME5* would serve as potential candidates for elucidating their precise roles in C_4_ photosynthesis as well as stress tolerance. This report will also serve as a blueprint for future studies on functionally characterizing the C_4_ pathway genes in other species to delineate their involvement in physiological as well as molecular processes associated with growth, development and stress response.

## Acknowledgments

Dr. Mehanathan Muthamilarasan acknowledges DST INSPIRE Faculty Award from the Department of Science & Technology, Ministry of Science & Technology, Government of India. Mr. Roshan K Singh is thankful to the Council of Scientific & Industrial Research, Ministry of Science & Technology, Government of India for the Research Fellowship. The authors are thankful to DBT-eLibrary Consortium (DeLCON) for providing access to the e-resources.

## Author’s contributions

Conception and design: MP and MM; experimental analyses: MM, RKS, BVS and PD; interpretation of data: MM, NKS and MP; writing, review and revision of manuscript: MM; study supervision: MP and MM; all authors have read and approved the final manuscript

## Supplementary Information

**Supplementary Fig. S1.** Structural features of C_4_ photosynthetic genes identified in *Setaria italica* and *S. viridis*

**Supplementary Table S1.** List of primers used in the present study

**Supplementary Table S2.** Profile of C_4_ photosynthetic genes identified in *Setaria italica* and *S. viridis*

**Supplementary Table S3.** Domain composition of architecture of C_4_ photosynthetic enzymes identified in *Setaria italica* and *S. viridis*

**Supplementary Table S4.** Summary of different *cis*-acting elements present in the upstream region of C_4_ photosynthetic genes

**Supplementary Table S5.** The Ka/Ks ratios and estimated divergence time of orthologous gene-pairs between *S. italica* and *P. virgatum*, *S. bicolor*, *Z. mays*, *B. distachyon* and *O. Sativa*

