## Supplemental Table S1 for "Molecular characterization and differential expression reveal functional divergence of stress-responsive enzymes in C_4_ panicoid models, *Setaria italica* and *Setaria viridis*"

**Supplementary Table S1.** List of primers used in the present study

| **Gene** | **Forward (5՛-3՛)** | **Reverse (5՛-3՛)** |
| --- | --- | --- |
| **qRT-PCR analysis** | | |
| *SiαCaH1* | GCCAGACCCTTTCCTTTTCC | CTTGCAGCCCTCCGAGTATAA |
| *SiγCaH3* | GCTGACGGCTATGAGGACAAC | GGCGGAAGATGCTGATGTCT |
| *SiβCaH5* | TGCACGACGACATGGAGAAG | AGGCTGACGTCTCTGCTGTTC |
| *SiPEPC1* | GGAGGACTCGGCCTTTAAGAA | AGGGATTTGTAGCACAGCTCAAG |
| *SiPEPC2* | CGCTCCGTGCAATTCCAT | CCAAGCCAAACAGGGAGATG |
| *SiPEPC-b* | CCCCGCGAGAACCTTACC | ACGCTCGCAGGTGTTGTACA |
| *SiPPDK1* | GGTCGCAAAGCTAGGCCTAA | GAAGGCTCCCCACCATGTT |
| *SiPPDK2* | TTGCTGCGGGCCTACCT | CTGCAAAGACGATCTGACCTACA |
| *SiMDH2* | TCTATCGTGCAGGGTCTTCATG | TCGCCGAAAGGTCCATCTT |
| *SiMDH3* | TCTGCCCTCGAAGCTCATG | TTGCTGGGTTGGCAACAAC |
| *SiMDH5* | TGAGCGCTGTGGTGCAAA | GGCAAAGTCCTCAAACTGAATGA |
| *SiMDH8* | GCACTGCCCAAACGCTCTT | TGCAATTGGGACAGTGGAGTT |
| *SiNADP-ME5* | GTGGGAGAGGTTCTTGGACTTG | AGCATGCCCACCAATGACA |
| *SiNADP-ME6* | GCCACCCGTGTCCATGAG | TGGCCTGATCAGCTAGTGCTT |
| *SiACT2* | CGCATATGTGGCTCTTGACT | GGGCACCTAAATCTCTCTGC |
| **Cloning for yeast overexpression** | | |
| *SiNADP-ME5* | GTGGGAGAGGTTCTTGGACTTG | AGCATGCCCACCAATGACA |
