## Supplemental Table S5 for "Molecular characterization and differential expression reveal functional divergence of stress-responsive enzymes in C_4_ panicoid models, *Setaria italica* and *Setaria viridis*"

**Supplementary Table S5.** The Ka/Ks ratios and estimated divergence time of orthologous gene-pairs between *S. italica* and *P. virgatum*, *S. bicolor*, *Z. mays*, *B. distachyon* and *O. Sativa*.

| ***S. italica – P. virgatum*** | | | | | | | | | | | | |
| --- | --- | --- | --- | --- | --- | --- | --- | --- | --- | --- | --- | --- |
| **Gene** | **Phytozome ID** | **Chr.** | **Start** | **End** | **Pv gene ID** | **Chr.** | **Start** | **End** | **Ka** | **Ks** | **Ka/Ks** | **Time of divergence (mya)** |
| SiαCaH1 | Seita.1G181000 | 1 | 26068546 | 26074416 | Pavir.1KG318900 | Chr01K | 56646428 | 56654835 | 0.06 | 0.04 | 1.47 | 3.3 |
| SiαCaH2 | Seita.2G228200 | 2 | 33024840 | 33026375 | Pavir.6NG274100 | Chr06N | 69972201 | 69973942 | 0.01 | 0.05 | 0.19 | 4.1 |
| SiαCaH4 | Seita.4G219300 | 4 | 33907822 | 33909573 | Pavir.2KG356500 | Chr02K | 68540642 | 68546910 | 0.05 | 0.06 | 0.89 | 4.4 |
| SiβCaH6 | Seita.5G240100 | 5 | 30342283 | 30345472 | Pavir.5NG263700 | Chr05N | 73104229 | 73108228 | 0.05 | 0.04 | 1.22 | 3.2 |
| SiβCaH7 | Seita.5G240200 | 5 | 30350751 | 30354837 | Pavir.5KG453400 | Chr05K | 78813778 | 78817399 | 0.04 | 0.06 | 0.61 | 4.8 |
| SiαCaH8 | Seita.6G179900 | 6 | 30373646 | 30375321 | Pavir.6KG306300 | Chr06K | 60505385 | 60506941 | 0.01 | 0.06 | 0.14 | 4.9 |
| SiαCaH12 | Seita.7G090500 | 7 | 19435708 | 19438028 | Pavir.7NG173400 | Chr07N | 43377868 | 43380931 | 0.01 | 0.08 | 0.14 | 6.4 |
| SiαCaH13 | Seita.7G325400 | 7 | 35467106 | 35467640 | Pavir.3KG031500 | Chr03K | 2339408 | 2340618 | 0.03 | 0.08 | 0.37 | 6.0 |
| SiαCaH14 | Seita.8G034800 | 8 | 2257456 | 2258363 | Pavir.3NG018000 | Chr03N | 1450358 | 1452072 | 0.03 | 0.08 | 0.38 | 5.8 |
| SiPEPC1 | Seita.1G020700 | 1 | 1784608 | 1791259 | Pavir.1NG132600 | Chr01N | 16673314 | 16679919 | 0.03 | 0.09 | 0.33 | 6.8 |
| SiPEPC2 | Seita.2G173700 | 2 | 26198697 | 26204510 | Pavir.2KG207600 | Chr02K | 33530574 | 33537066 | 0.02 | 0.04 | 0.4 | 3.3 |
| SiPEPC5 | Seita.5G324500 | 5 | 37380598 | 37387338 | Pavir.5KG551100 | Chr05K | 94932624 | 94939109 | 0.02 | 0.05 | 0.35 | 3.8 |
| SiPPDK1 | Seita.3G247900 | 3 | 21259472 | 21273920 | Pavir.3KG261700 | Chr03K | 45028963 | 45044628 | 0.03 | 0.05 | 0.54 | 4.0 |
| SiPPDK2 | Seita.9G354600 | 9 | 41401305 | 41409346 | Pavir.9KG370100 | Chr09K | 58645465 | 58654342 | 0.02 | 0.05 | 0.38 | 3.5 |
| SiMDH1 | Seita.2G137100 | 2 | 15714933 | 15719067 | Pavir.6KG413200 | Chr06K | 72200618 | 72206502 | 0.02 | 0.06 | 0.26 | 5.0 |
| SiMDH2 | Seita.2G401000 | 2 | 46370023 | 46372520 | Pavir.2NG596600 | Chr02N | 98004734 | 98007114 | 0.02 | 0.06 | 0.26 | 4.7 |
| SiMDH3 | Seita.3G137700 | 3 | 9555248 | 9559625 | Pavir.3KG214200 | Chr03K | 19006879 | 19011797 | 0.02 | 0.07 | 0.22 | 5.5 |
| SiMDH4 | Seita.3G276100 | 3 | 25100252 | 25102599 | Pavir.6NG143300 | Chr06N | 57694395 | 57697295 | 0.01 | 0.04 | 0.35 | 3.2 |
| SiMDH5 | Seita.3G401800 | 3 | 50147307 | 50151954 | Pavir.3KG562000 | Chr03K | 71284993 | 71289755 | 0.01 | 0.09 | 0.1 | 6.8 |
| SiMDH9 | Seita.7G189800 | 7 | 26571855 | 26574500 | Pavir.7NG273600 | Chr07N | 57519622 | 57522277 | 0.04 | 0.08 | 0.45 | 6.5 |
| SiMDH10 | Seita.9G073100 | 9 | 4282933 | 4286388 | Pavir.9NG074500 | Chr09N | 4973629 | 4976825 | 0.01 | 0.04 | 0.2 | 3.2 |
| SiMDH11 | Seita.9G222200 | 9 | 16519383 | 16524072 | Pavir.9NG301600 | Chr09N | 32711085 | 32715216 | 0.08 | 0.04 | 1.89 | 3.4 |
| SiNADP-ME1 | Seita.2G322000 | 2 | 41047172 | 41055500 | Pavir.2KG446000 | Chr02K | 78913666 | 78922157 | 0.08 | 0.05 | 1.77 | 3.7 |
| SiNADP-ME2 | Seita.3G109300 | 3 | 7278989 | 7282388 | Pavir.3KG194100 | Chr03K | 17251687 | 17255473 | 0.13 | 0.04 | 3.03 | 3.2 |
| SiNADP-ME3 | Seita.3G284800 | 3 | 26806748 | 26808875 | Pavir.5NG039100 | Chr05N | 6652456 | 6657648 | 0.13 | 0.05 | 2.3 | 4.2 |
| SiNADP-ME6 | Seita.5G301800 | 5 | 35587449 | 35592805 | Pavir.5KG515900 | Chr05K | 92078529 | 92083750 | 0.13 | 0.06 | 2.21 | 4.4 |
| SiNADP-ME7 | Seita.5G314300 | 5 | 36545367 | 36550031 | Pavir.5KG546100 | Chr05K | 94459137 | 94463852 | 0.13 | 0.06 | 2.05 | 4.7 |
| SiNADP-ME9 | Seita.9G200600 | 9 | 14379420 | 14385175 | Pavir.8KG331900 | Chr08K | 67892437 | 67898578 | 0.02 | 0.07 | 0.24 | 5.0 |
| **Mean** | | | | | | | | | **0.04** | **0.06** | **0.81** | **4.56** |
| ***S. italica – S. bicolor*** | | | | | | | | | | | | |
| **Gene** | **Phytozome ID** | **Chr.** | **Start** | **End** | **Sb gene ID** | **Chr.** | **Start** | **End** | **Ka** | **Ks** | **Ka/Ks** | **Time of divergence (mya)** |
| SiαCaH1 | Seita.1G181000 | 1 | 26068546 | 26074416 | Sobic.004G166000 | Sb4 | 51559903 | 51563175 | 0.07 | 0.61 | 0.12 | 47.2 |
| SiαCaH2 | Seita.2G228200 | 2 | 33024840 | 33026375 | Sobic.002G224000 | Sb2 | 61530646 | 61535739 | 0.04 | 0.27 | 0.16 | 21.1 |
| SiαCaH4 | Seita.4G219300 | 4 | 33907822 | 33909573 | Sobic.010G189366 | Sb10 | 53000019 | 53001351 | 0.05 | 0.31 | 0.17 | 24.1 |
| SiβCaH7 | Seita.5G240200 | 5 | 30350751 | 30354837 | Sobic.003G234600 | Sb3 | 57323907 | 57327962 | 0.08 | 0.35 | 0.24 | 26.8 |
| SiαCaH9 | Seita.6G180000 | 6 | 30395216 | 30397511 | Sobic.007G155100 | Sb7 | 58862663 | 58864411 | 0.02 | 0.33 | 0.07 | 25.4 |
| SiPEPC1 | Seita.1G020700 | 1 | 1784608 | 1791259 | Sobic.004G106900 | Sb4 | 10201753 | 10208601 | 0.05 | 0.17 | 0.26 | 13.4 |
| SiPEPC2 | Seita.2G173700 | 2 | 26198697 | 26204510 | Sobic.002G167000 | Sb2 | 52122128 | 52127601 | 0.05 | 0.35 | 0.15 | 26.7 |
| SiPEPC3 | Seita.4G175200 | 4 | 28034663 | 28043265 | Sobic.004G106900 | Sb4 | 10201753 | 10208601 | 0.07 | 0.61 | 0.12 | 46.9 |
| SiPEPC4 | Seita.5G147000 | 5 | 13061164 | 13065839 | Sobic.007G106500 | Sb7 | 38045106 | 38050518 | 0.05 | 0.27 | 0.16 | 21.1 |
| SiPEPC5 | Seita.5G324500 | 5 | 37380598 | 37387338 | Sobic.003G301800 | Sb3 | 63252659 | 63260241 | 0.08 | 0.48 | 0.18 | 36.6 |
| SiPPDK1 | Seita.3G247900 | 3 | 21259472 | 21273920 | Sobic.009G132900 | Sb9 | 48726109 | 48738853 | 0.08 | 0.28 | 0.28 | 21.3 |
| SiPPDK2 | Seita.9G354600 | 9 | 41401305 | 41409346 | Sobic.001G326900 | Sb1 | 61372602 | 61380996 | 0.09 | 0.29 | 0.32 | 22.1 |
| SiMDH1 | Seita.2G137100 | 2 | 15714933 | 15719067 | Sobic.007G166300 | Sb7 | 60149510 | 60153216 | 0.06 | 0.35 | 0.16 | 26.7 |
| SiMDH2 | Seita.2G401000 | 2 | 46370023 | 46372520 | Sobic.002G385700 | Sb2 | 74038066 | 74041773 | 0.06 | 0.32 | 0.18 | 24.5 |
| SiMDH3 | Seita.3G137700 | 3 | 9555248 | 9559625 | Sobic.009G240700 | Sb9 | 57799754 | 57804273 | 0.09 | 0.23 | 0.39 | 18.0 |
| SiMDH4 | Seita.3G276100 | 3 | 25100252 | 25102599 | Sobic.007G137600 | Sb7 | 56567941 | 56570679 | 0.04 | 0.19 | 0.2 | 14.6 |
| SiMDH6 | Seita.5G245200 | 5 | 30734735 | 30738108 | Sobic.003G238500 | Sb3 | 57798790 | 57804310 | 0.09 | 0.19 | 0.44 | 15.0 |
| SiMDH7 | Seita.6G159300 | 6 | 28219985 | 28221178 | Sobic.007G137600 | Sb7 | 56567941 | 56570679 | 0.06 | 0.53 | 0.12 | 40.9 |
| SiMDH8 | Seita.6G251800 | 6 | 35757958 | 35761505 | Sobic.007G166300 | Sb7 | 60149510 | 60153216 | 0.05 | 0.21 | 0.22 | 16.0 |
| SiMDH9 | Seita.7G189800 | 7 | 26571855 | 26574500 | Sobic.006G170800 | Sb6 | 52734301 | 52736982 | 0.03 | 0.43 | 0.07 | 33.2 |
| SiMDH11 | Seita.9G222200 | 9 | 16519383 | 16524072 | Sobic.001G219300 | Sb1 | 20443589 | 20448787 | 0.07 | 0.71 | 0.1 | 54.6 |
| SiNADP-ME5 | Seita.5G134300 | 5 | 11688238 | 11693473 | Sobic.009G108700 | Sb9 | 43545770 | 43551420 | 0.09 | 0.25 | 0.38 | 19.0 |
| SiNADP-ME6 | Seita.5G301800 | 5 | 30330923 | 30338082 | Sobic.003G234200 | Sb3 | 57297987 | 57310502 | 0.04 | 0.51 | 0.07 | 39.1 |
| SiNADP-ME7 | Seita.5G314300 | 5 | 36545367 | 36550031 | Sobic.003G292400 | Sb3 | 62488685 | 62493225 | 0.06 | 0.35 | 0.16 | 26.7 |
| SiNADP-ME9 | Seita.9G200600 | 9 | 14379420 | 14385175 | Sobic.001G201700 | Sb1 | 18302412 | 18308518 | 0.06 | 0.32 | 0.18 | 24.5 |
| **Mean** | | | | | | | | | **0.06** | **0.36** | **0.20** | **27.42** |
| ***S. italica – Zea mays*** | | | | | | | | | | | | |
| **Gene** | **Phytozome ID** | **Chr.** | **Start** | **End** | **Zm gene ID** | **Chr.** | **Start** | **End** | **Ka** | **Ks** | **Ka/Ks** | **Time of divergence (mya)** |
| SiαCaH1 | Seita.1G181000 | 1 | 26068546 | 26074416 | GRMZM5G807267 | Zm2 | 57684823 | 57686916 | 0.11 | 0.26 | 0.43 | 20.3 |
| SiαCaH2 | Seita.2G228200 | 2 | 33024840 | 33026375 | GRMZM2G088208 | Zm2 | 185673128 | 185674652 | 0.06 | 0.37 | 0.16 | 28.4 |
| SiαCaH4 | Seita.4G219300 | 4 | 33907822 | 33909573 | GRMZM2G009633 | Zm9 | 103408327 | 103410418 | 0.07 | 0.28 | 0.26 | 21.2 |
| SiβCaH7 | Seita.5G240200 | 5 | 30350751 | 30354837 | GRMZM2G121878 | Zm3 | 215547876 | 215555228 | 0.05 | 0.37 | 0.15 | 28.3 |
| SiαCaH9 | Seita.6G180000 | 6 | 30395216 | 30397511 | GRMZM2G113165 | Zm1 | 199139161 | 199142195 | 0.04 | 0.34 | 0.12 | 26.2 |
| SiPEPC1 | Seita.1G020700 | 1 | 1784608 | 1791259 | GRMZM2G074122 | Zm4 | 227810824 | 227818618 | 0.04 | 0.5 | 0.07 | 38.2 |
| SiPEPC2 | Seita.2G173700 | 2 | 26198697 | 26204510 | GRMZM2G473001 | Zm7 | 86459173 | 86464913 | 0.07 | 0.37 | 0.18 | 28.4 |
| SiPEPC3 | Seita.4G175200 | 4 | 28034663 | 28043265 | GRMZM2G083841 | Zm9 | 62306266 | 62311673 | 0.06 | 0.59 | 0.1 | 45.4 |
| SiPEPC4 | Seita.5G147000 | 5 | 13061164 | 13065839 | GRMZM2G074122 | Zm4 | 227810824 | 227818618 | 0.07 | 0.51 | 0.15 | 38.9 |
| SiPEPC5 | Seita.5G324500 | 5 | 37380598 | 37387338 | GRMZM2G110714 | Zm8 | 173256053 | 173268872 | 0.05 | 0.33 | 0.17 | 25.6 |
| SiPPDK1 | Seita.3G247900 | 3 | 21259472 | 21273920 | GRMZM2G097457 | Zm8 | 106530187 | 106541052 | 0.03 | 0.22 | 0.15 | 17.1 |
| SiPPDK2 | Seita.9G354600 | 9 | 41401305 | 41409346 | GRMZM2G306345 | Zm6 | 146179867 | 146189965 | 0.08 | 0.32 | 0.25 | 25.0 |
| SiMDH1 | Seita.2G137100 | 2 | 15714933 | 15719067 | GRMZM2G129513 | Zm1 | 203209824 | 203213983 | 0.08 | 0.7 | 0.12 | 54.2 |
| SiMDH2 | Seita.2G401000 | 2 | 46370023 | 46372520 | GRMZM2G141289 | Zm7 | 167677886 | 167680080 | 0.03 | 0.35 | 0.07 | 26.7 |
| SiMDH3 | Seita.3G137700 | 3 | 9555248 | 9559625 | GRMZM2G154595 | Zm6 | 165831537 | 165836297 | 0.06 | 0.29 | 0.2 | 22.2 |
| SiMDH5 | Seita.3G401800 | 3 | 50147307 | 50151954 | GRMZM2G072744 | Zm1 | 282605048 | 282608688 | 0.03 | 0.34 | 0.08 | 26.1 |
| SiMDH6 | Seita.5G245200 | 5 | 30734735 | 30738108 | GRMZM2G466833 | Zm3 | 213978038 | 213986724 | 0.06 | 0.37 | 0.16 | 28.4 |
| SiMDH7 | Seita.6G159300 | 6 | 28219985 | 28221178 | GRMZM2G161245 | Zm1 | 213144280 | 213146855 | 0.06 | 0.32 | 0.18 | 24.3 |
| SiMDH9 | Seita.7G189800 | 7 | 26571855 | 26574500 | GRMZM2G101290 | Zm2 | 20538918 | 20540574 | 0.11 | 0.46 | 0.23 | 35.7 |
| SiMDH10 | Seita.9G073100 | 9 | 4282933 | 4286388 | GRMZM2G072744 | Zm1 | 282605048 | 282608688 | 0.11 | 0.48 | 0.22 | 36.6 |
| SiNADP-ME3 | Seita.3G284800 | 3 | 26806748 | 26808875 | GRMZM2G085019 | Zm3 | 7276387 | 7281737 | 0.08 | 0.42 | 0.2 | 32.5 |
| SiNADP-ME5 | Seita.5G134300 | 5 | 11688238 | 11693473 | GRMZM2G122479 | Zm6 | 139464390 | 139470075 | 0.09 | 0.37 | 0.25 | 28.4 |
| SiNADP-ME6 | Seita.5G301800 | 5 | 35587449 | 35592805 | GRMZM2G159724 | Zm3 | 201756871 | 201761835 | 0.05 | 0.28 | 0.18 | 21.4 |
| SiNADP-ME7 | Seita.5G314300 | 5 | 36545367 | 36550031 | GRMZM2G118770 | Zm8 | 174612565 | 174617106 | 0.06 | 0.29 | 0.2 | 22.7 |
| SiNADP-ME8 | Seita.7G040900 | 7 | 12423964 | 12426409 | GRMZM2G085019 | Zm3 | 7276387 | 7281737 | 0.06 | 0.26 | 0.22 | 19.9 |
| SiNADP-ME9 | Seita.9G200600 | 9 | 14379420 | 14385175 | GRMZM2G085747 | Zm5 | 23916399 | 23923301 | 0.07 | 0.24 | 0.28 | 18.3 |
| **Mean** | | | | | | | | | **0.06** | **0.37** | **0.18** | **28.48** |
| ***S. italica – O. sativa*** | | | | | | | | | | | | |
| **Gene** | **Phytozome ID** | **Chr.** | **Start** | **End** | **Os gene ID** | **Chr.** | **Start** | **End** | **Ka** | **Ks** | **Ka/Ks** | **Time of divergence (mya)** |
| SiαCaH1 | Seita.1G181000 | 1 | 26068546 | 26074416 | LOC_Os02g33030 | Os2 | 19636271 | 19638854 | 0.3 | 0.69 | 0.40 | 49.3 |
| SiβCaH7 | Seita.5G240200 | 5 | 30350751 | 30354837 | LOC_Os01g45274 | Os1 | 25700370 | 25705090 | 0.04 | 1.20 | 0.03 | 44.21 |
| SiPEPC1 | Seita.1G020700 | 1 | 1784608 | 1791259 | LOC_Os02g14770 | Os2 | 8177490 | 8184220 | 0.10 | 0.57 | 0.18 | 31.50 |
| SiPEPC2 | Seita.2G173700 | 2 | 26198697 | 26204510 | LOC_Os09g14670 | Os9 | 8692191 | 8697573 | 0.06 | 0.69 | 0.09 | 53.12 |
| SiPEPC3 | Seita.4G175200 | 4 | 28034663 | 28043265 | LOC_Os01g11054 | Os2 | 8177490 | 8184220 | 0.06 | 0.43 | 0.15 | 39.95 |
| SiPEPC4 | Seita.5G147000 | 5 | 13061164 | 13065839 | LOC_Os05g33570 | Os1 | 5899555 | 5909595 | 0.3 | 0.69 | 0.40 | 49.3 |
| SiPPDK1 | Seita.3G247900 | 3 | 21259472 | 21273920 | LOC_Os03g31750 | Os5 | 19718506 | 19737857 | 0.06 | 0.69 | 0.09 | 53.12 |
| SiPPDK2 | Seita.9G354600 | 9 | 41401305 | 41409346 | LOC_Os08g44810 | Os3 | 18153143 | 18160127 | 0.10 | 0.45 | 0.22 | 35.42 |
| SiMDH1 | Seita.2G137100 | 2 | 15714933 | 15719067 | LOC_Os01g46070 | Os8 | 28141042 | 28146270 | 0.10 | 0.46 | 0.21 | 39.67 |
| SiMDH3 | Seita.3G137700 | 3 | 9555248 | 9559625 | LOC_Os08g33720 | Os1 | 26190752 | 26194517 | 0.3 | 0.69 | 0.40 | 49.3 |
| SiMDH6 | Seita.5G245200 | 5 | 30734735 | 30738108 | LOC_Os01g09320 | Os1 | 26190752 | 26194517 | 0.3 | 0.69 | 0.40 | 49.3 |
| SiMDH7 | Seita.6G159300 | 6 | 28219985 | 28221178 | LOC_Os01g54030 | Os8 | 21054659 | 21057561 | 0.26 | 0.71 | 0.40 | 50.7 |
| SiMDH9 | Seita.7G189800 | 7 | 26571855 | 26574500 | LOC_Os04g46560 | Os4 | 27605166 | 27608347 | 0.3 | 0.69 | 0.40 | 49.3 |
| SiMDH10 | Seita.9G073100 | 9 | 4282933 | 4286388 | LOC_Os03g56280 | Os3 | 32086001 | 32089685 | 0.06 | 0.69 | 0.09 | 53.12 |
| SiNADP-ME3 | Seita.3G284800 | 3 | 26806748 | 26808875 | LOC_Os10g35960 | Os1 | 4738905 | 4744591 | 0.3 | 0.69 | 0.40 | 49.3 |
| SiNADP-ME5 | Seita.5G134300 | 5 | 11688238 | 11693473 | LOC_Os10g35960 | Os1 | 4738905 | 4744591 | 0.06 | 0.69 | 0.09 | 53.12 |
| SiNADP-ME7 | Seita.5G314300 | 5 | 36545367 | 36550031 | LOC_Os01g54030 | Os1 | 31076571 | 31081377 | 0.10 | 0.45 | 0.22 | 35.42 |
| SiNADP-ME9 | Seita.9G200600 | 9 | 14379420 | 14385175 | LOC_Os10g35960 | Os10 | 19218567 | 19224021 | 0.10 | 0.46 | 0.21 | 39.67 |
| **Mean** | | | | | | | | | **0.16** | **0.65** | **0.24** | **45.82** |
| ***S. italica – B. distachyon*** | | | | | | | | | | | | |
| **Gene** | **Phytozome ID** | **Chr.** | **Start** | **End** | **Bd gene ID** | **Chr.** | **Start** | **End** | **Ka** | **Ks** | **Ka/Ks** | **Time of divergence (mya)** |
| SiαCaH1 | Seita.1G181000 | 1 | 26068546 | 26074416 | Bradi3g44940 | Bd3 | 46634866 | 46638114 | 0.06 | 0.41 | 0.14 | 43.13 |
| SiαCaH2 | Seita.2G228200 | 2 | 33024840 | 33026375 | Bradi3g38260 | Bd3 | 40544556 | 40546768 | 0.26 | 0.52 | 0.50 | 37.1 |
| SiαCaH4 | Seita.4G219300 | 4 | 33907822 | 33909573 | Bradi1g36190 | Bd1 | 31991500 | 31992734 | 0.3 | 0.69 | 0.40 | 49.3 |
| SiαCaH9 | Seita.6G180000 | 6 | 30395216 | 30397511 | Bradi3g38260 | Bd3 | 40544556 | 40546768 | 0.3 | 0.87 | 0.30 | 62.1 |
| SiPEPC1 | Seita.1G020700 | 1 | 1784608 | 1791259 | Bradi3g09210 | Bd3 | 7351388 | 7358870 | 0.26 | 0.71 | 0.40 | 50.7 |
| SiPEPC4 | Seita.5G147000 | 5 | 13061164 | 13065839 | Bradi2g06620 | Bd2 | 5068548 | 5072711 | 0.26 | 0.52 | 0.50 | 37.1 |
| SiPPDK1 | Seita.3G247900 | 3 | 21259472 | 21273920 | Bradi2g25745 | Bd2 | 23900805 | 23916710 | 0.3 | 0.69 | 0.40 | 49.3 |
| SiMDH1 | Seita.2G137100 | 2 | 15714933 | 15719067 | Bradi3g12460 | Bd3 | 11175330 | 11180410 | 0.26 | 0.83 | 0.30 | 59.3 |
| SiMDH3 | Seita.3G137700 | 3 | 9555248 | 9559625 | Bradi2g45200 | Bd2 | 45357840 | 45361976 | 0.28 | 0.75 | 0.40 | 53.6 |
| SiMDH7 | Seita.6G159300 | 6 | 28219985 | 28221178 | Bradi3g37140 | Bd3 | 39196420 | 39198805 | 0.3 | 0.87 | 0.30 | 62.1 |
| SiMDH9 | Seita.7G189800 | 7 | 26571855 | 26574500 | Bradi5g17700 | Bd5 | 21056008 | 21058735 | 0.26 | 0.52 | 0.50 | 37.1 |
| SiMDH10 | Seita.9G073100 | 9 | 4282933 | 4286388 | Bradi1g07170 | Bd1 | 4995776 | 4999732 | 0.3 | 0.69 | 0.40 | 49.3 |
| SiNADP-ME3 | Seita.3G284800 | 3 | 26806748 | 26808875 | Bradi2g05620 | Bd2 | 4100048 | 4105539 | 0.3 | 0.69 | 0.40 | 49.3 |
| SiNADP-ME5 | Seita.5G134300 | 5 | 11688238 | 11693473 | Bradi2g05620 | Bd2 | 4100048 | 4105539 | 0.06 | 0.41 | 0.14 | 43.13 |
| SiNADP-ME6 | Seita.5G301800 | 5 | 35587449 | 35592805 | Bradi2g48570 | Bd2 | 48596284 | 48602026 | 0.3 | 0.87 | 0.30 | 62.1 |
| **Mean** | | | | | | | | | **0.25** | **0.67** | **0.36** | **49.64** |
