## Supplementary figures and images for "Molecular characterization and differential expression reveal functional divergence of stress-responsive enzymes in C_4_ panicoid models, *Setaria italica* and *Setaria viridis*"

### Supplemental Figure S1

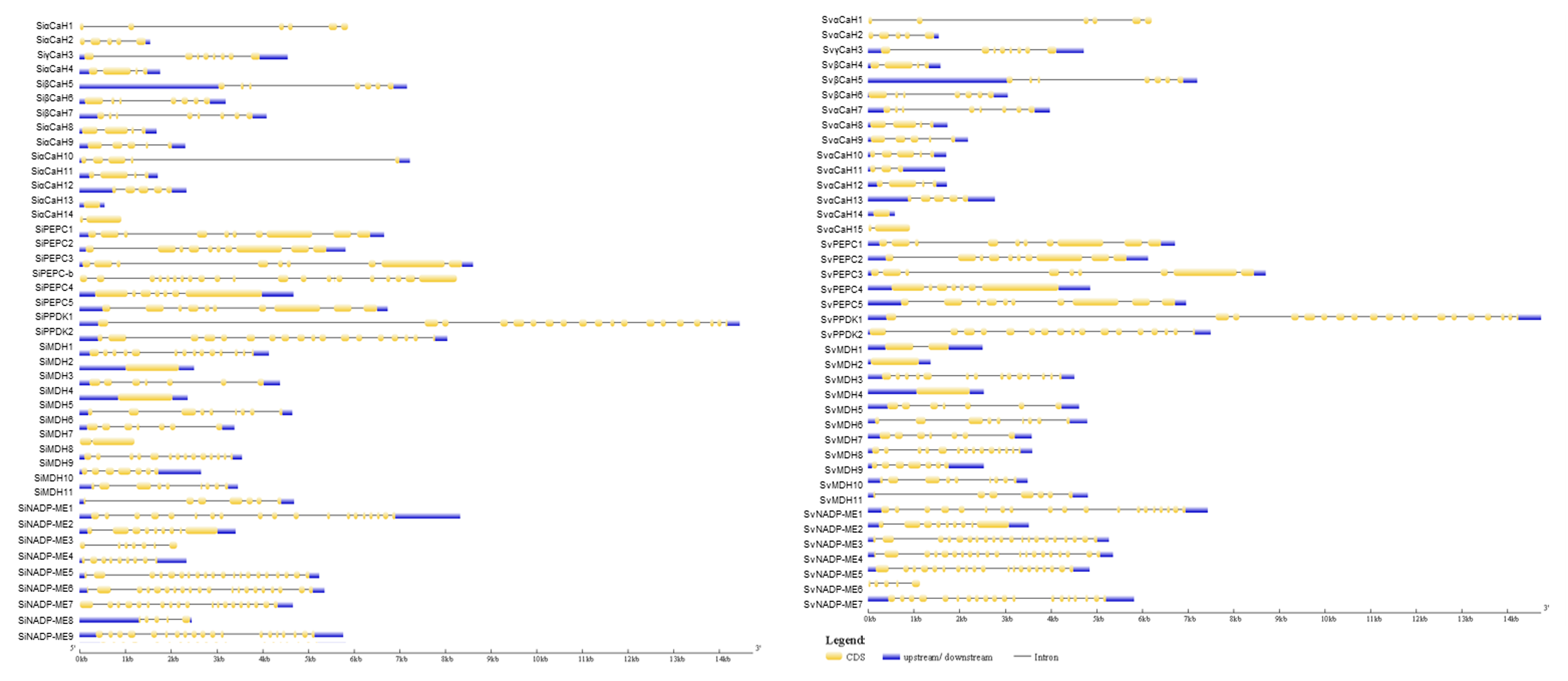
